# NADPH-oxidase 2 is required for molecular adaptations to high-intensity interval training in skeletal muscle

**DOI:** 10.1101/542514

**Authors:** Carlos Henríquez-Olguín, Leila Baghersad Renani, Lyne Arab-Ceschia, Steffen H. Raun, Aakash Bhatia, Zhencheng Li, Jonas R. Knudsen, Rikard Holmdahl, Thomas E. Jensen

## Abstract

**Objective:** Reactive oxygen species (ROS) have been proposed as signaling molecules mediating exercise training adaptation, but the ROS source has remained unclear. This study aimed to investigate the requirement for NADPH oxidase (NOX)2-dependent redox changes induced by acute and long-term high-intensity interval training (HIIT) in skeletal muscle in a mouse model lacking functional NOX2 complex due to deficient p47phox (*Ncf1*) subunit expression (*ncf1\** mutation).

**Methods:** HIIT was investigated after an acute bout of exercise and after a chronic intervention (3x week for 6 weeks) in wildtype (WT) vs. NOX2 activity-deficient (*ncf1\**) mice. NOX2 activation during HIIT was measured using a genetically-encoded biosensor. Immunoblotting and single-fiber staining were performed to measure classical exercise-training responsive endpoints in skeletal muscle.

**Results:** A single bout of HIIT increased NOX2 activity measured using electroporated p47roGFP oxidation immediately after exercise but not 1h after exercise. After a 6-week of HIIT regime, improvements in maximal running capacity and some muscle training-markers responded less to HIIT in the *ncf1\** mice compared to WT, including superoxide dismutase (SOD)2, catalase, hexokinase II (HK II), pyruvate dehydrogenase (PDH) and protein markers of mitochondrial oxidative phosphorylation complexes. Strikingly, HIIT-training increased mitochondrial network area and decreased fragmentation in WT mice only.

**Conclusion:** This study provided evidence that HIIT exercise activates NOX2 complex in skeletal muscle and that the presence of functional NOX2 is required for specific skeletal muscle adaptations to HIIT relating to antioxidant defense, glucose metabolism, and mitochondria.

GRAPHICAL ABSTRACT

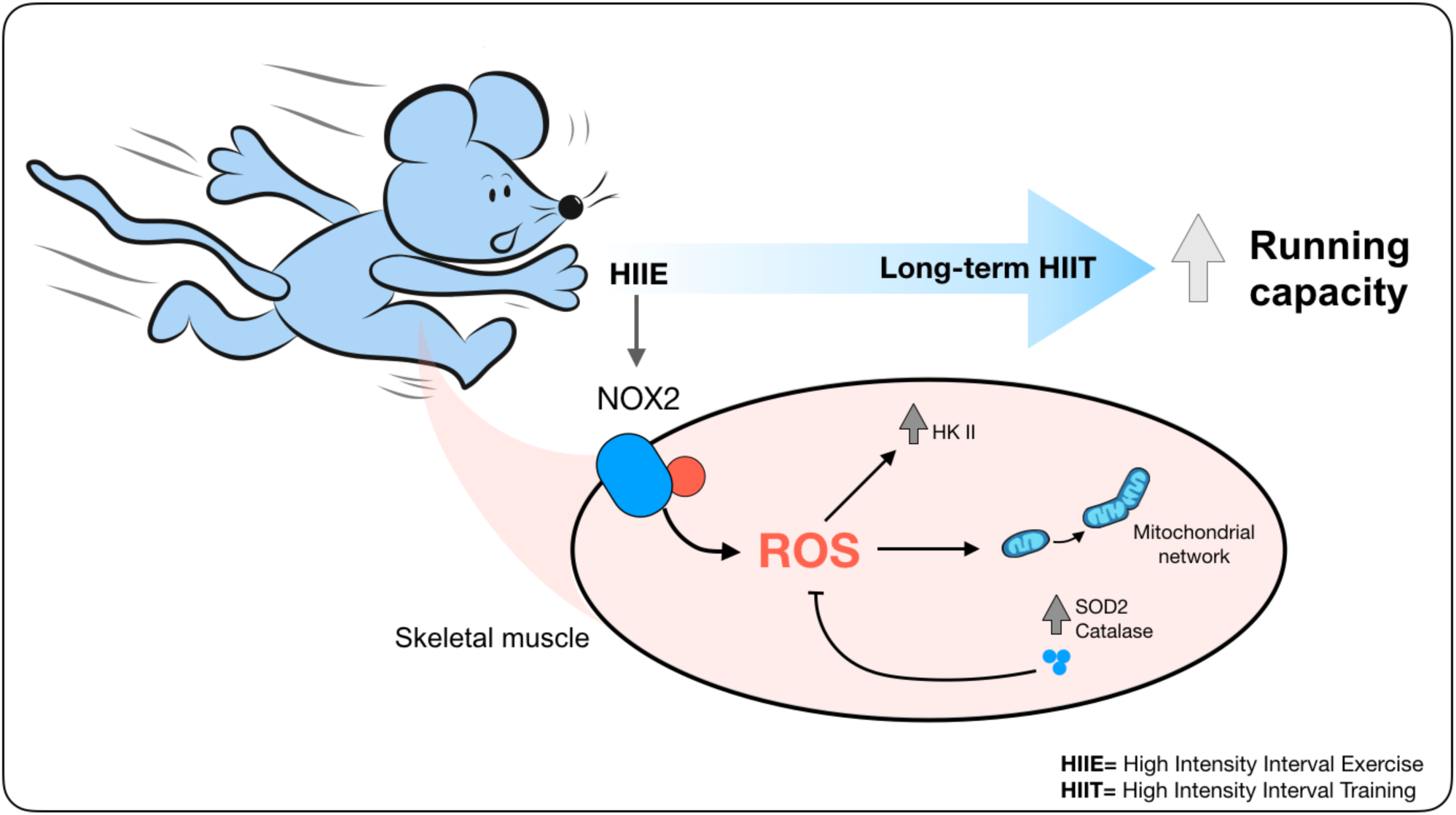

**Highlights:**

- Acute HIIE induces transient NOX2 complex activity *in vivo* in muscle
- Skeletal muscle adaptations to HIIT were impaired in *ncf1*-deficient mice
- Functional NOX2 is necessary for HIIT-induced increased expression of antioxidants enzymes
- *Ncf1*-deficient mice lack HIIT-induced mitochondrial adaptations

## 1. INTRODUCTION

Physical inactivity is regarded as cause of morbidity and premature mortality worldwide [1]. Inactivity and sedentary behavior are estimated to be responsible for between 6%-10% of the burden of disease from non-communicable diseases [2]. Exercise intensity is a significant variable explaining the health benefits induced by physical activity [3, 4]. Indeed, structured high-intensity interval training (HIIT) has been demonstrated to improve both whole-body and skeletal muscle metabolic health in different populations [5-7]. Despite the proven efficacy of HIIT to promote metabolic health, the underlying mechanisms improving adaptation in the HIIT-exercised musculature are not yet fully understood. Gaining a deeper understanding of the mechanistic basis of the signaling mechanisms governing the acute and chronic responses to HIIT in skeletal muscle would support the development of more effective exercise training regimes and the identification of potential drug targets.

Reactive oxygen species (ROS) act as intracellular compartmentalized second messengers mediating skeletal muscle adaptations in both health and disease [8, 9]. Sprint interval bicycle exercise has been shown to elicit greater post-exercise plasma hydrogen peroxide compared to moderate exercise in humans [10], suggesting that exercise-induced ROS production in skeletal muscle may be intensity dependent. Specific ROS may be required for adaptation to chronic exercise, since ROS scavengers have been shown to disrupt some of the acute and long-term responses to exercise in skeletal muscle (reviewed in [11]). Furthermore, elevated levels of systemic oxidative stress markers were associated with greater adaptations after 6-weeks of exercise training in humans [12]. Taken together, this suggests that ROS may contribute to the intensity-dependent myocellular exercise training adaptation in skeletal muscle.

Although many studies have suggested the importance of ROS molecules to exercise training adaptation, the exact myocellular source of ROS has remained unclear [13]. For many years, mitochondria were believed to be as the primary source of ROS during exercise in skeletal muscle. More recently, non-mitochondrial sources have emerged as potential ROS sources during contractile activity in skeletal muscle [14, 15]. Based on studies using electrically evoked contractions in isolated rodent muscle, the professional superoxide-producing enzyme complex NADPH oxidase 2 (NOX2) was strongly suggested to mediate contraction-induced ROS production [15]. Moreover, pharmacological inhibition of NOX2 has been shown to disrupt acute signaling and gene expression elicited by moderate-intensity endurance exercise in mice [14]. Thus, whether NOX2 is a significant source of ROS during high-intensity exercise and its role in the specific context of HIIT requires clarification.

In the present study, we investigated whether NOX2 is activated in mouse skeletal muscle by physiological acute high-intensity exercise (HIIE). Furthermore, we investigated if NOX2 activity was required for the long-term skeletal muscle adaptations to HIIT using a mouse model lacking functional NOX2 due to a mutation in its regulatory subunit p47phox. We hypothesized that NOX2 was a major ROS source in skeletal muscle during this high-intensity exercise modality and required for long-term HIIT adaptations.

## 2. RESULTS

### 2.1 Acute HIIE increased NOX2-dependent redox changes in skeletal muscle

To assess NOX2-specific ROS production, we used a genetically-encoded probe expressing human p47^phox^ fused to the N-terminus of redox-sensitive green fluorescent protein 2 (p47roGFP) [16]. Oxidation of roGFP in this probe causes a ratio metric change in fluorescence, measureable as a reduction in the 470 nm and an increase/maintenance in the 405 nm emission. The p47roGFP reporter was expressed in tibialis anterior muscle (TA) via *in vivo* electroporation 1 week before exercise. Acute HIIE elicited an increase in p47roGFP oxidation immediately after exercise (time 0) which returned to baseline at 1h after HIIE (Figure 1). This demonstrated that NOX2-dependent ROS production is increased transiently by HIIE.

**Figure 1.**
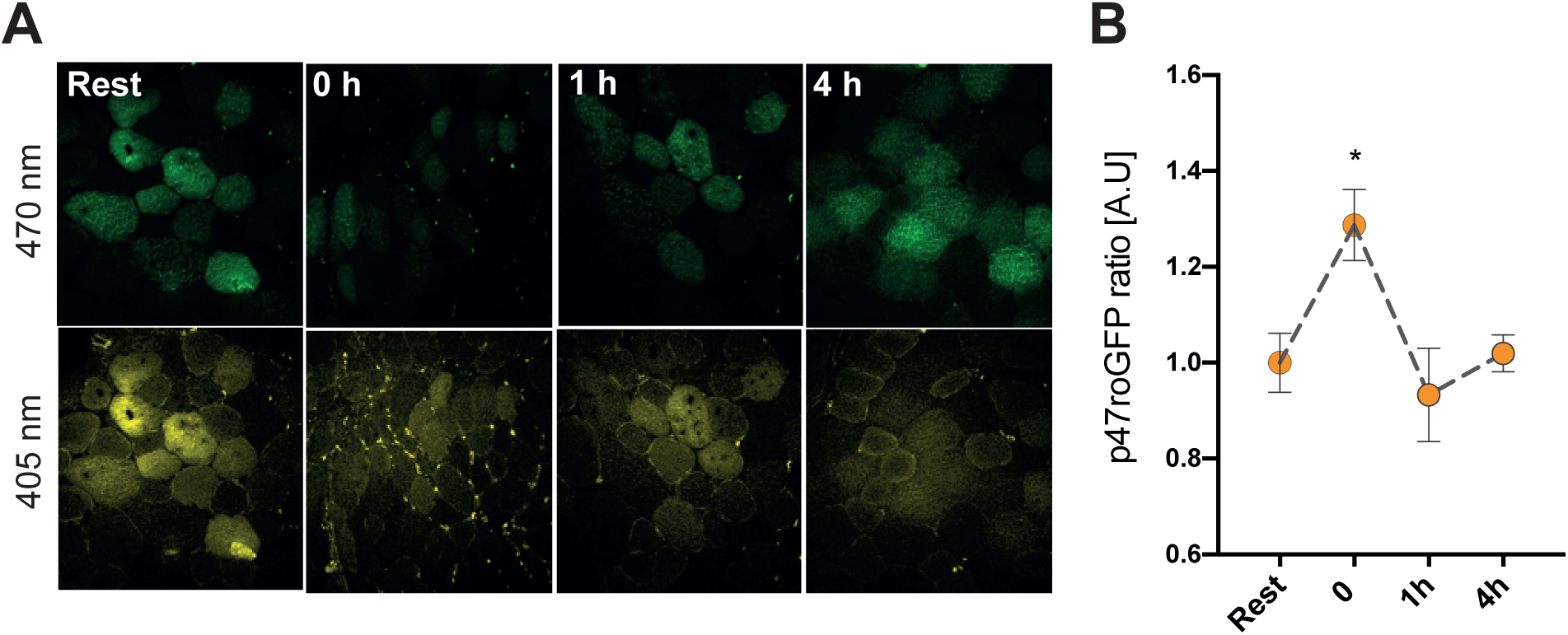
High-intensity interval exercise activates NOX2 transiently in skeletal muscle. A) Representative images and b) ratio metric quantification (405/470 nm) of p47roGFP biosensor signal at resting, exercise and post-exercise period in tibialis anterior muscle. * denotes p< 0.05 compared to the resting condition using a one-way ANOVA followed by a Holm-Sidak’s multiple comparisons test. Values are mean ± SEM (n=4 per group).

Phosphorylation of p38 MAPK and ERK1/2 were increased by acute HIIE (Figure 2). Moreover, phosphorylation of AMPK and its substrate ACC2 increased immediately after exercise in Quad and SOL muscles (Figure 2).

**Figure 2.**
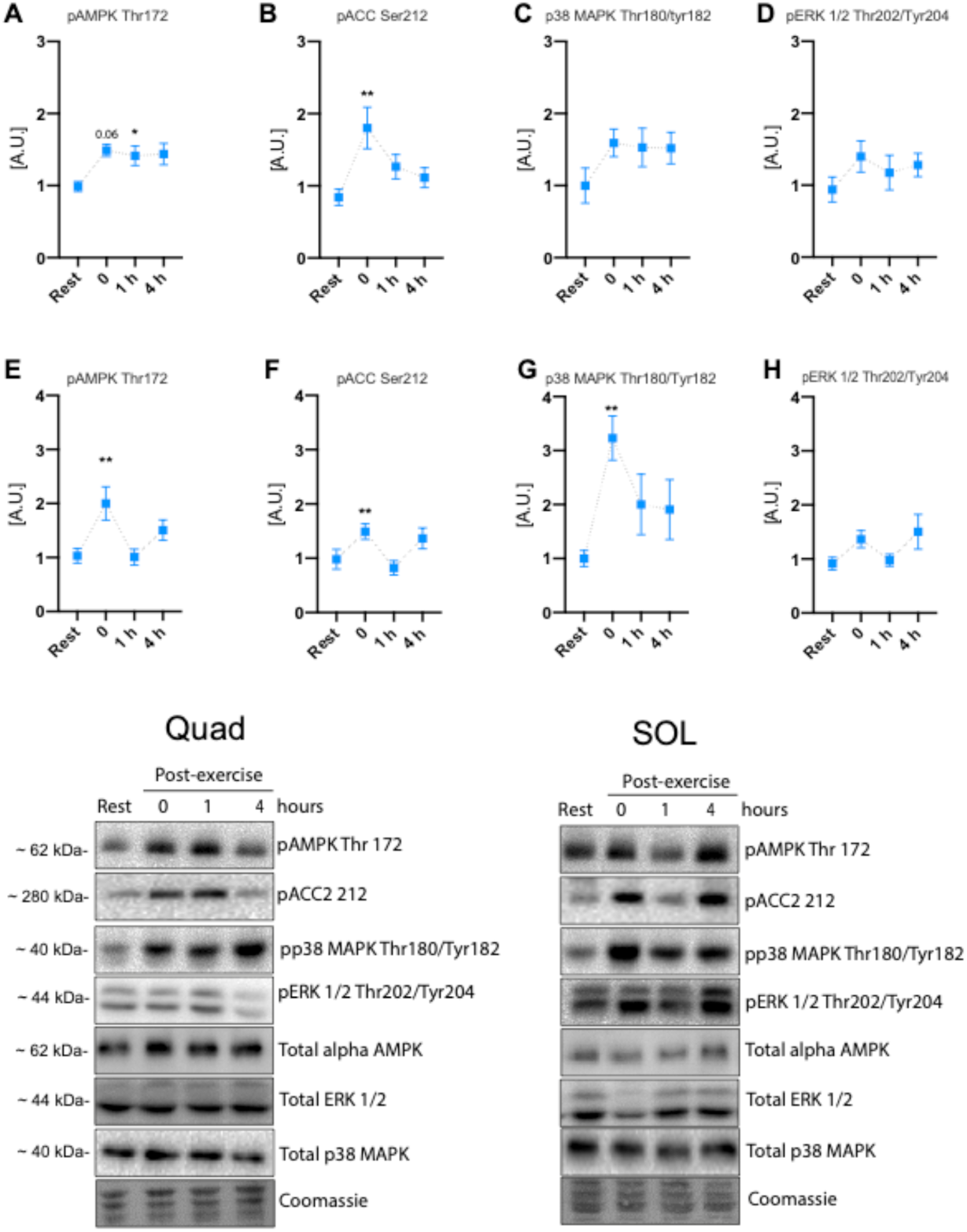
Acute cellular signaling induced by high-intensity interval exercise. Exercise-stimulated phosphorylation of A) pAMPK Thr172 B) pACC Ser212 C) p38 MAPK Thr180/Tyr182 D) pERK 1/2 Thr202/Tyr204 for quadriceps muscle and E) pAMPK Thr172 F) pACCC Ser212 G) p38 MAPK Thr180/Tyr182 H) pERK 1/2 Thr202/Tyr204 for soleus muscle. ^*/**^denotes p< 0.05 and p< 0.01, respectively, compared to the resting condition using one-way ANOVA followed by a Holm-Sidak’s multiple comparisons test. Values are mean ± SEM (n=8 per group).

To evaluate whether an acute HIIE increased the autophagy-associated LC3 I lipidation in skeletal muscle, we measured LC3 I and II expression in muscle lysates. Neither Quad nor SOL muscle changed their LC3 II/I ratio up to 4h post-HIIT (Figure 3), suggesting that HIIE does not increase autophagy during exercise or in the immediate post-exercise recovery period.

**Figure 3.**
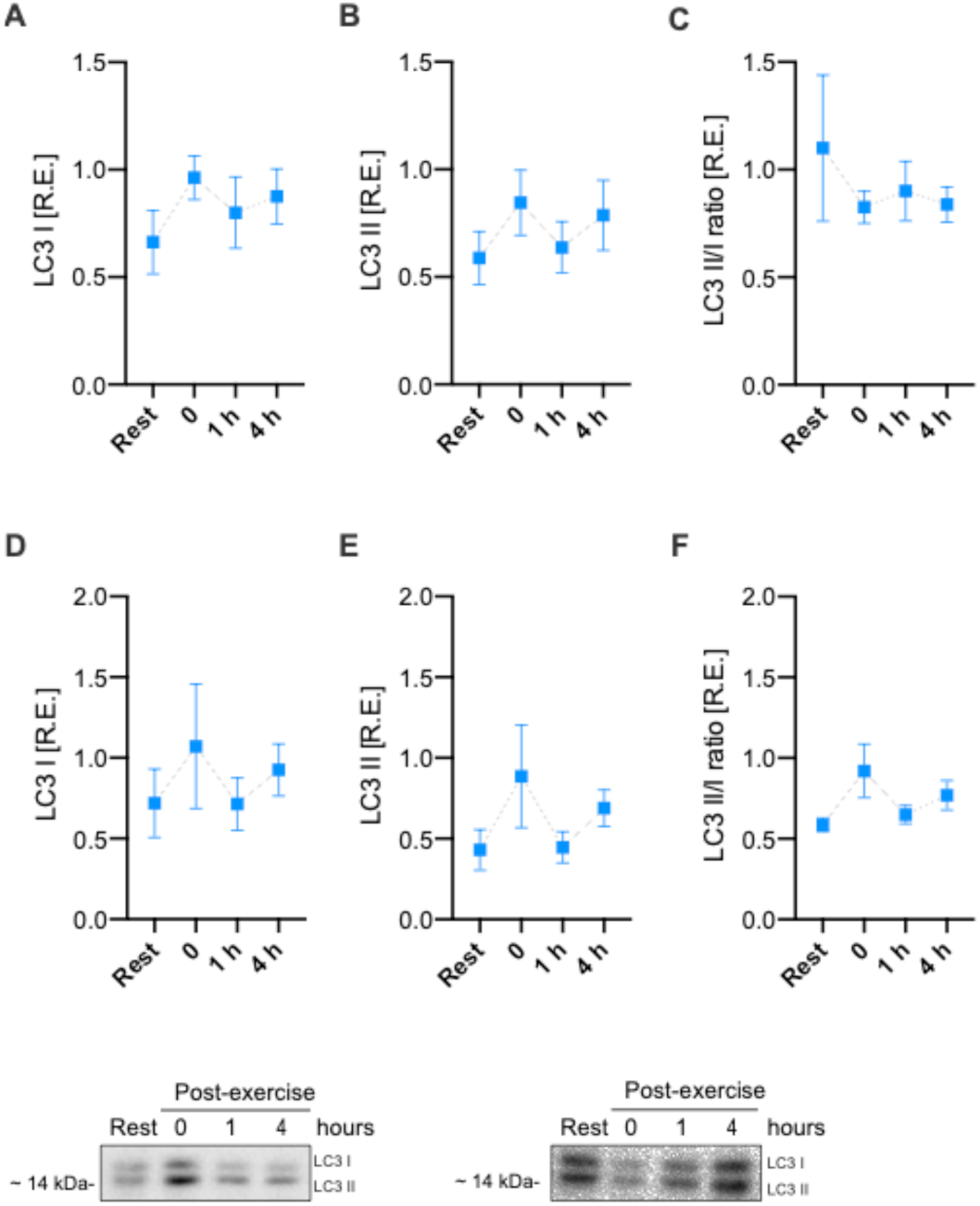
Autophagy markers are not induced by acute high-intensity interval exercise. Exercise-stimulated changes in LC3-I, LC3-II, and LC3-II/LC3-I ratio. One-way ANOVA was performed. Values are mean ± SEM (n=8 per group).

### 2.2 Lack of NOX2 complex activity reduced responsiveness in running capacity

Having confirmed that an acute bout of HIIE activated NOX2, we next tested whether NOX2 is required for long-term HIIT adaptation. We used *Ncf1\** mice, a described whole-body loss-of-function model for NOX2 activity [17] with undetectable ROS production [18]. *Ncf1\** mice were indistinguishable from WT and no cases of the previously reported spontaneous post-partum arthritis[19] were observed during >1y of breeding. Parallel results from our laboratory demonstrated the absence of in vivo treadmill exercise-stimulated ROS production in skeletal muscle of *Ncf1\** mice [20], in agreement with earlier in vitro studies using electrical stimulation [16].

Presently, WT and *ncf1\** mice performed a three-times per week HIIT regimen for 6-weeks on a motorized treadmill [21]. The HIIT intervention increased maximal running capacity in WT but not *ncf1\** mice, compared to untrained mice (Figure 4A: +23% vs. 10% in WT and *ncf1\** respectively). Similar body weight and composition were observed in the untrained state, but trained *ncf1\** mice displayed lower body weight compared to trained WT mice (Figure 4B). Reduced body fat and increased lean mass were observed in the *ncf1\** HIIE group compared with the trained WT group (Figure 4C-D). Neither training nor genotype affected energy intake (Figure 4 E-F). Overall, *ncf1\** mice showed a tendency towards lower RER than WT mice (Figure 4G), but a significant genotype main effect was only observed in the trained group (Figure 4H). Similar oxygen consumption was observed between genotypes in both the untrained and trained state (Figure 4I-J). Thus, NOX2-deficient mice display impaired HIIT-induced improvements in maximal running capacity and lower body fat content after HIIT training.

**Figure 4.**
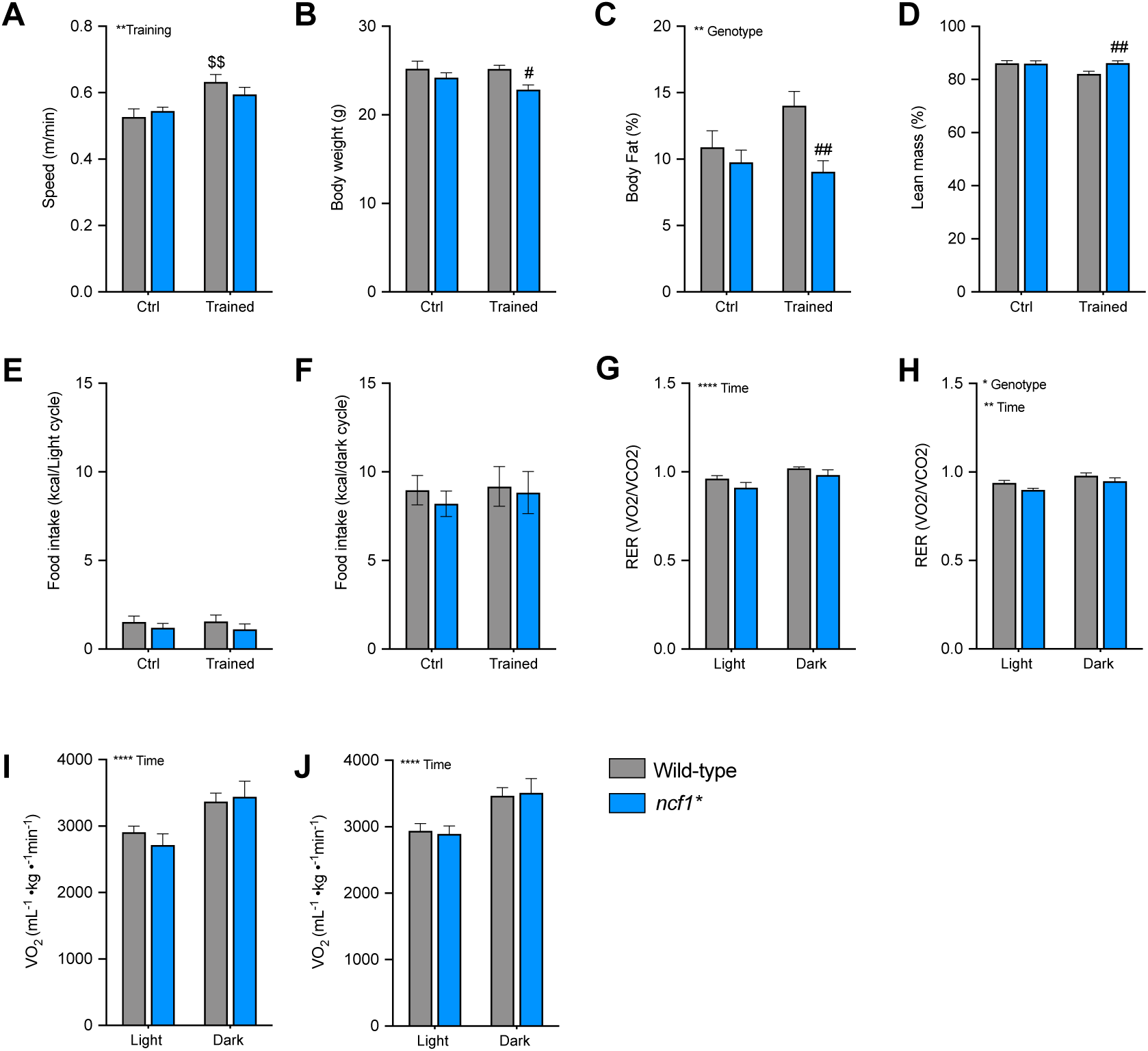
*ncf1\** mice show a lower responsiveness in running capacity after long-term HIIT. 6-weeks high-intensity interval training outcomes in A) maximal running speed B) Body weight C-D) Body composition E) Food intake during the light and F) dark cycle Respiratory exchange ratio in G) untrained and H) trained mice, oxygen uptake in I) Untrained and J) trained mice. ^$$^ denotes p<0.01 vs. WT Ctrl group, ^##^ denotes p<0.01 vs. WT trained mice. Two-way ANOVA was performed to test for effects of training, genotype, and time, followed by a Holm-Sidak’s *post hoc* test corrected for multiple comparisons. Values are mean ± SEM (n=7-8).

### 2.3 NOX2 deficiency impaired the HIIT-induced increase in specific antioxidant enzymes

ROS scavengers decrease the moderate-intensity training-stimulated improvements in antioxidant capacity in skeletal muscle [22, 23]. Furthermore, acute pharmacological blockade of NOX2 reduces mRNA levels of key antioxidant enzymes in mouse skeletal muscle [14]. Here, the mitochondrial-localized manganese-dependent superoxide dismutase (SOD2) protein expression increased only in WT (+119%) but not in *ncf1\** (+26%) in response to HIIT, driven by a higher baseline SOD2 content in *ncf1\** vs. WT mice (Figure 5A). A similar tendency was observed for catalase expression in WT (+74%, p = 0.06) vs. *ncf1\** (26%) (Figure 5B). Neither genotype nor exercise training affected thioredoxin reductase 2 (TRX2), gp91phox, or nNOS protein expression (Figure 5C-E). Thus, NOX2 deficiency reduces the HIIT adaptations in specific antioxidant enzymes but not globally in redox-signaling proteins.

**Figure 5.**
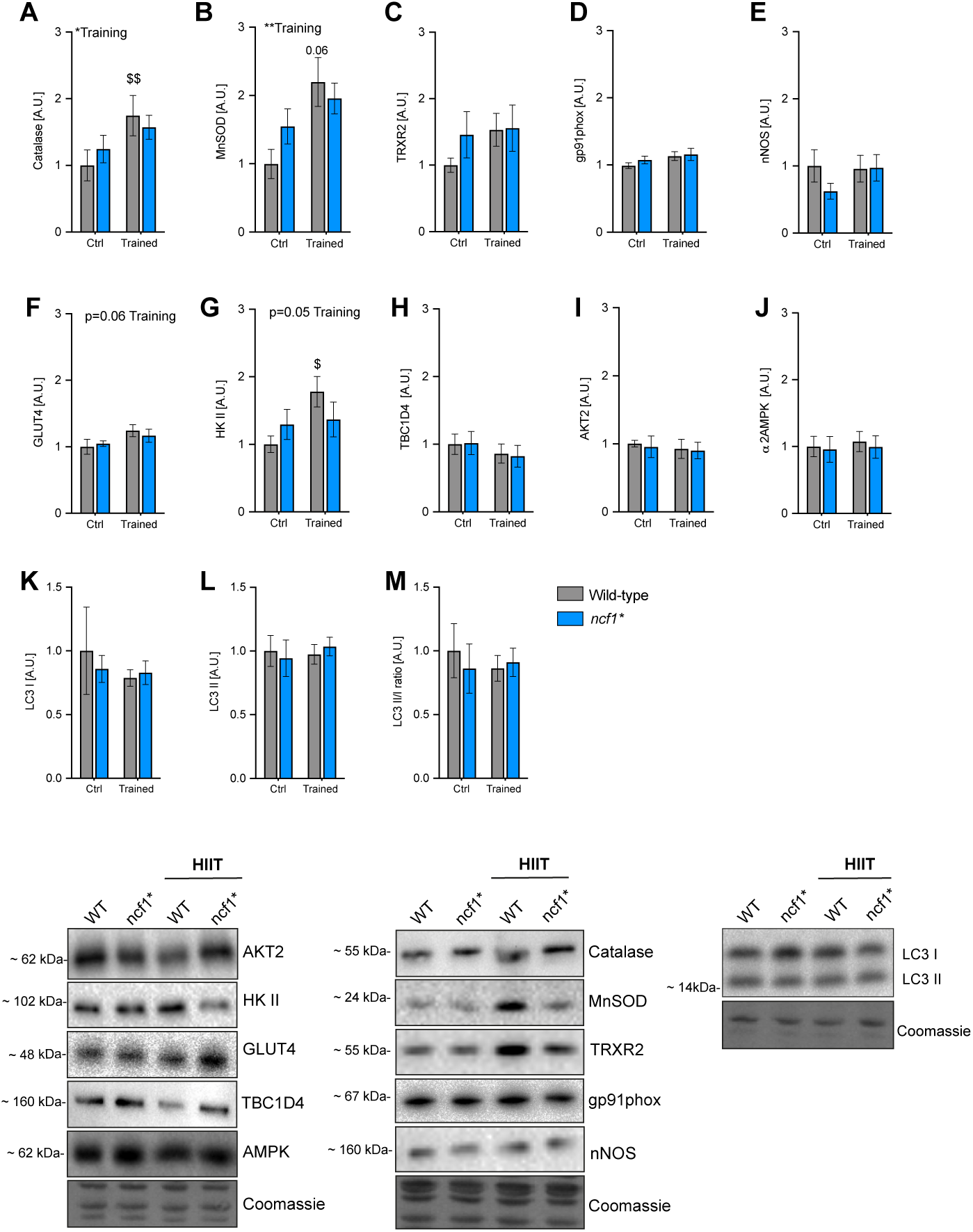
Multiple exercise-training markers in *ncf1\** muscles are less responsive to 6-weeks of HIIT. A-E) High intensity interval training-induced changes in redox-related proteins, F-J) Glucose handling related proteins and K-M) autophagy-related proteins. ^$, $$^ denotes p< 0.05 and p< 0.01, respectively compared to WT ctrl Two-way ANOVA was performed to test for effects of training and genotype, followed by a Holm-Sidak’s *post hoc* test corrected for multiple comparisons. Values are mean ± SEM (n=7-8).

### 2.4 HIIT-induced muscle HKII expression was NOX2-dependent

Exercise training may improve muscle insulin sensitivity partially by increasing the protein content of glucose handling proteins [24]. Since antioxidants decrease exercise training-induced insulin sensitivity [22], we tested whether HIIT and/or NOX2 deficiency affected glucose handling proteins. Neither HIIT nor genotype affected glucose transport 4 (GLUT4) expression (Figure 5F). In contrast, hexokinase II, a rate-limiting glycolytic enzyme, increased +77% with HIIT in WT (p<0.05) but did not respond to HIIT in the *ncf1\** mice (Figure 5G). No effects were observed on total TBCD4 or AKT2 protein abundance (Figure 5H-I).

### 2.5 Lack of NOX2 activity impaired mitochondrial adaptations to HIIT

HIIT has been associated with an increase in both mitochondrial content and function [25]. ROS could regulate training-induced mitochondrial biogenesis [26, 27]. Moreover, pharmacological inhibition of NOX2 blunts exercise-stimulated mRNA levels of mitochondrial enzymes [14]. To test whether NOX2 activity is required for training-induced mitochondrial biogenesis, we first measured mitochondrial electron transport chain complex protein abundance and other mitochondrial proteins. The mitochondrial complex I marker was increased (+55%) in the trained group compared to the sedentary control in WT but not significantly increased in *ncf1\** mice (Figure 6A). A genotype main effect was found for complex III and Complex IV markers (Figure 6C-D). Moreover, total PDH levels increased in the WT (+129%) but not in *ncf1\** mice (+30%, Figure 6G).

**Figure 6.**
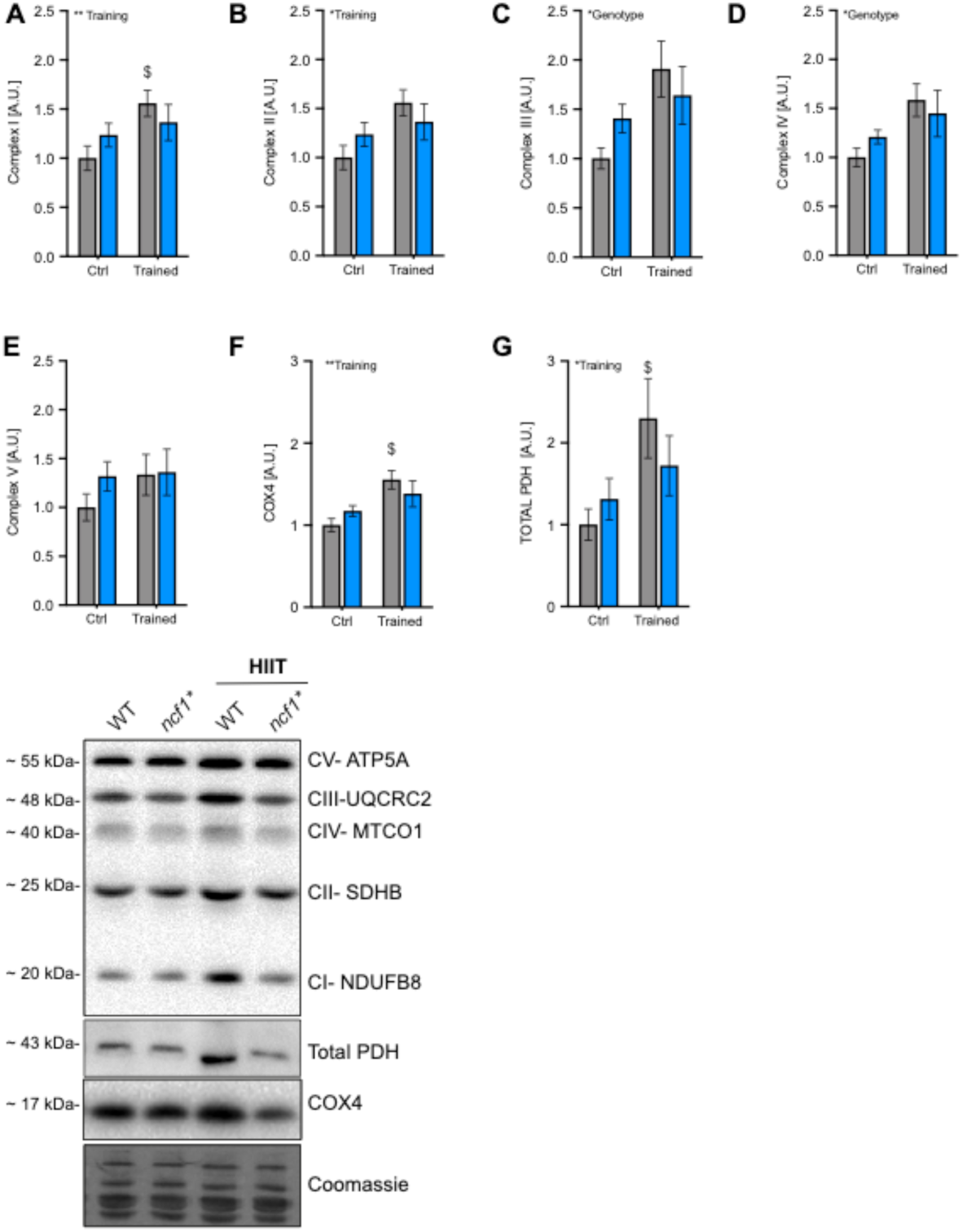
Mitochondrial-related proteins increase to a smaller extent after HIIT in mice lacking functional NOX2 activity. Mitochondrial electron chain complexes A) Complex I B) Complex II C) Complex III D) Complex IV B) Complex V F) COX4 and G) PDH. ^$^ denotes p<0.05 vs. WT Ctrl group. Two-way ANOVA was performed to test for effects of training, and genotype, followed by a Holm-Sidak’s *post hoc* test corrected for multiple comparisons. Values are mean ± SEM (n=7-8).

Mitochondria in skeletal muscle undergo fission/fusion-events and can vary from highly fragmented to interconnected tubular networks [28]. Since exercise training has been associated with more elongated and fused mitochondria [29, 30], we estimated the change in the mitochondrial network in single quadriceps muscle fibers by immunofluorescent imaging of COX4 (Figure 7A-D). Mitochondrial fragmentation was reduced by HIIT training in WT but not *ncf1\** mice (Figure 7B), and the cytosolic area occupied by mitochondria was increased by HIIT in WT but not in *ncf1\** mice (Figure 7C). Consistent with a disturbed ability to fuse mitochondria in *ncf1\** mice, we observed an increase of the inner membrane fusion protein OPA1 by HIIT in WT but not in *ncf1\** mice, and a tendency towards the same for the mitochondrial outer membrane fusion protein mitofusin-2 (MFN2) (p=0.05) (Figure 7D-F). Taken together, this shows that NOX2 is required for multiple aspects of mitochondrial adaptations to HIIT, with particularly strong effects on mitochondrial network morphology.

**Figure 7.**
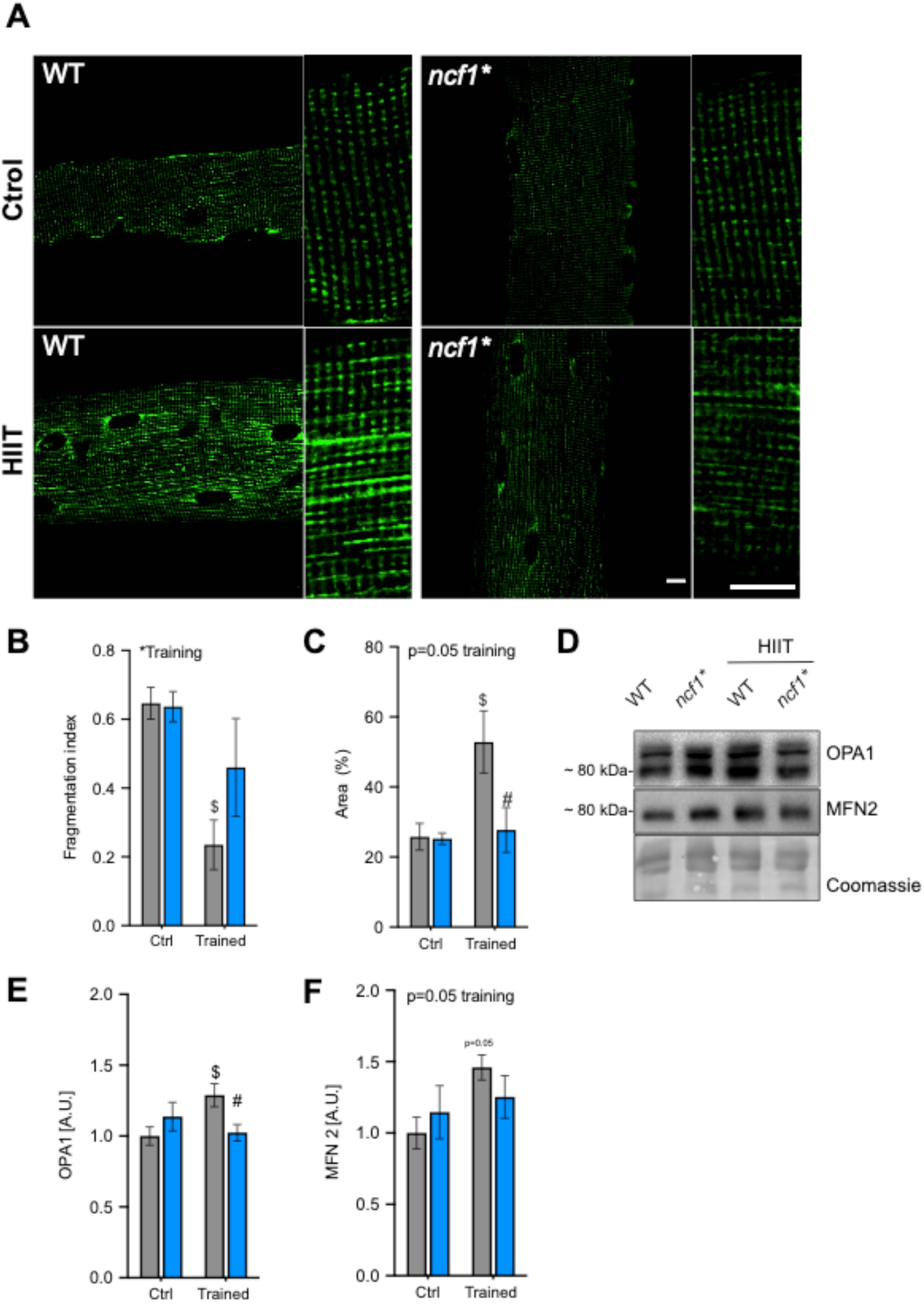
HIIT increases mitochondrial fragmentation and related proteins in skeletal muscle of wildtype mice only. **A)** Representative images from mitochondria-specific stains in quadriceps single fibers. Bars indicate 10 μm. B) Mitochondrial fragmentation index C) Mitochondrial area D) representative and quantification of E) OPA1 and F) MFN2 protein abundance. $ denotes p< 0.05, respectively compared to WT Ctrl. ^#^denotes p<0.05 WT trained vs. *ncf1\** trained mice Two-way ANOVA was performed to test for effects of training and genotype, followed by a Holm-Sidak’s *post hoc* test corrected for multiple comparisons. Values are mean ± SEM (A-C, n=4 and E-F, n=7-8).

## 3. DISCUSSION

ROS are proposed to act as signaling molecules mediating myocellular exercise training adaptations [9]. However, the exact source of ROS involved is still debated and may differ depending on the intensity of exercise. A recent study showed that adaptive gene expression to acute *in vivo* and *in vitro* exercise models was blunted by NOX2 inhibitors [14]. Our current study supports and extends on these data by showing that the functional improvement in maximal exercise capacity, a number of exercise-responsive proteins and most strikingly, mitochondrial morphological adaptations were less responsive to HIIT training in mice lacking functional NOX2 complex. The specific adaptations are briefly discussed below followed by some closing reflections on their potential connectivity.

### 3.1 Running performance

Some [31-33] but not all studies [23, 27] have found that ROS scavengers impair the training-induced improvements in maximal exercise capacity in rodents and humans. The lack of consistency in previous studies has been attributed to differences in antioxidant supplementation efficacy, specificity and/or training regimens. Here, using a genetic loss-of-function model, we observed that an acute bout of HIIE was sufficient to induce p47roGFP biosensor oxidation in skeletal muscle, indicating that HIIE activated NOX2 *in vivo*, and that lack of NOX2 activity in skeletal muscle was associated with a blunted improvement in running capacity after a 6-week HIIT period. An immediate concern is that the decreased running capacity in *ncf1\** vs. WT mice after HIIT could have contributed to the decreased protein response to HIIT. However, we find this unlikely since 1) there was no difference in running capacity before training and the mean difference in maximal running speed after training was only ∼5%, arguing that these mice were exposed to a similar HIIT intensity and volume and 2) some training-responsive proteins such as GLUT4 responded equally to HIIT in both genotypes. Overall, these data suggest that NOX2 is likely a major contributor to the previously proposed ROS-mediated increase in endurance exercise capacity in rodents and humans [11].

### 3.2 Antioxidant capacity

Chronic adaptations to exercise training are thought to result from the cumulative stimulatory effect of repeated acute exercise bouts on gene transcription [34]. Three weeks of HIIT in humans improve antioxidant capacity in plasma of humans [35]. This is consistent with the proposal that ROS are essential mediators of the hormetic increase in endogenous antioxidant defense in response to training [36]. A previous study found that NOX2 was required for some of the acute antioxidant defense gene transcription responses to muscle activity since pharmacological blockade of NOX2 *in vivo* or electrically stimulated contraction *in vitro* blocked exercise-responsive gene transcription of SOD2 and glutathione peroxidase transcription [14]. This was suggested to be regulated by a NOX2-dependent NF-κB pathway [37]. Whether NF-kB was a contributing factor for the lack of HIIT response in current study was not investigated.

### 3.3 Glucose handling enzymes

Both GLUT4 and HKII are known to respond to HIIT in humans [38]. A key finding in our study was that HKII, arguably one of the most exercise training-responsive proteins in man, did not increase after training in the NOX2-deficient *ncf1\** mice. A ROS-dependence is consistent with previous studies showing that antioxidants blunt electrically stimulated contraction-induced HKII mRNA and activity in muscle cells [39, 40] and training-induced HKII mRNA in mice [41]. In contrast, the increase in GLUT4 expression with HIIT did not appear to depend on NOX2 activity.

### 3.4 Mitochondrial protein expression

Endurance exercise training and HIIT are known to increase the protein content, volume, and function of skeletal muscle mitochondria [42]. Previous studies reported that the antioxidants Vitamin C and E reduced exercise training-induced mitochondrial biogenesis markers such as COX4 and CS, likely by downregulation of PCG1-alpha and TFAM in human and murine skeletal muscle [22, 23, 26, 31]. Interestingly, exercise-induced TFAM and citrate synthase mRNA levels in mouse skeletal muscle were also blocked by pharmacological inhibition of NOX2 [14]. Presently, we observed that the well-established mitochondrial content marker COX4 [43], as well as the mitochondrial metabolic capacity-markers PDH and mitochondrial complex I, increased after HIIT in WT but not *ncf1\** mice. We speculate that the general tendency towards increased base-line expression of mitochondria-related proteins in *ncf1\** vs. WT mice might increase resting fat oxidation, consistent with the observed lower RER and leaner phenotype in trained *ncf1\** vs. WT mice.

### 3.5 Mitochondrial network morphology

Endurance exercise training is associated with an elongated mitochondrial network [29] and decreased mitochondrial fragmentation [30] in the trained musculature in rodents and humans [44]. HIIT has been reported to increase the respiratory capacity of mitochondria [25, 45] but the HIIT-associated changes in mitochondrial morphology have, as far as we are aware, not been investigated in any species. Our data show for the first time that these responses also occur with HIIT. Furthermore, the HIIT-induced mitochondrial network remodeling exhibited the strongest quantitative impairment of any endpoint in mice lacking NOX2 activity. The genotype difference in mitochondrial network was accompanied by a blunted HIIT-induced increases of the mitochondrial fusion related proteins OPA1 and MFN2 in *ncf1\** vs WT muscle. We cannot dismiss the changes in fusion-related proteins are independent of the changes in mitochondrial content. This might provide clues to the underlying mechanism but requires further investigation.

### 3.7. Are the different molecular findings connected?

Many of the proteins that differed significantly between WT and *ncf1\** mice displayed a similar expression pattern with an increased relative expression in untrained *ncf1\** mice and a lower relative responsiveness to HIIT in *ncf1\** mice. Importantly, the changes were confined to subsets of proteins and not observed for e.g. GLUT4, TBC1D4 or Akt2 expression. Based on the literature many of these changes could be mechanistically connected. Thus, the mitochondrial adaptations to HIIT were impaired both in terms of network morphology and mitochondrial protein expression. The baseline upregulation of several antioxidant enzymes, including SOD2 and catalase could suggest increased mitochondrial ROS production. Mitochondrial ROS production has previously been associated with mitochondrial fragmentation and higher resting fat oxidation [46-48], but their contribution here is unclear since genotype differences in these parameters were only observed in the trained state. Worth noting, however, we did observe a significant decrease in baseline RER in *ncf1\** vs. WT mice [20], indicating increased resting fat oxidation in *ncf1\** mice. Hexokinase II is known to shuttle to and from mitochondria [49]. Interestingly, acute disruption of HKII binding to mitochondria, using a peptide-inhibitor in a perfused mouse heart ischemia-reperfusion injury model, markedly reduced cardiac recovery and increased ROS during ischemia and reperfusion [50]. Similar links have been established in other tissues, including skeletal muscle [51]. Thus, we speculate that HKII may somehow be connected to both the increased base-line fat oxidation and changes in mitochondrial morphology/function. The mechanistic connections between the observed molecular changes in *ncf1\** vs WT muscle should be measured in the future.

### 3.8 Conclusion

This study provided evidence that NOX2 is activated by HIIE and that the presence of functional NOX2 is required for long-term training adaptation, including increased muscle protein expression of antioxidant defense enzymes, mitochondrial enzymes and hexokinase II, and increased mitochondrial network volume and decreased mitochondrial fragmentation. This suggests that NOX2 signaling modulates the exercise-response to HIIT. If and how these changes are interconnected should be clarified in future studies.

## MATERIALS AND METHODS

### Animals

10-week-old female B10.Q wildtype (WT) and B10.Q. p47phox mutated (*ncf1*)* were maintained on a 12 hours light/dark cycle, group-housed with free access to water and standard rodent chow diet (Altromin no. 1324; Chr. Pedersen, Denmark).

All experiments were approved by the Danish Animal Experimental Inspectorate (2015–15–0201–00477).

### Maximal running capacity

Running capacity was carried as previously described [52]. Briefly, mice were acclimated to the treadmill three times (10 min at 0.16 m/s) (Treadmill TSE Systems) a week before the maximal running tests. The maximal running test started at 0.16 m/s for 300 s with 10 % incline, followed by a step-wise increase (0.2 m/s every 60s) in running speed until exhaustion. Exhaustion was defined as the point at which instead of running on the treadmill, the mice fall back on the grid three times within 30 secs. Maximal running speed was determined as the last completed stage during the incremental test. All tests were performed blinded.

### Indirect calorimetry and body composition

Body composition was assessed by MRI-scanning (EchoMRI-4, Echo Medical System LLC, Texas, USA) according to the manufacturer’s instructions. Whole-body metabolism was assessed in a 16-chamber indirect calorimetry system after a 2-day acclimation period (PhenoMaster; TSE Systems, Frankfurt, Germany).

### Acute high-intensity interval exercise (HIIE)

Fed mice performed a single HIIE bout switching between 2 min running at 100% of the maximal running speed of each group, and then 2 min of active recovery running at 30% of the maximal running speed for a total of 60 min. Tissues were harvest immediately, one and four hours after exercise.

### HIIT intervention

Both WT and *ncf1\** mice were randomly assigned to the control or HIIT group. The HIIT training was carried as previously described [21]. Briefly, HIIT training involved treadmill running three days per week for six weeks. In each training session, mice switched between two min running at 100% of the maximal running speed of each group, and then 2 min of active recovery running at 30% of the maximal running speed for a total of 60 min. During the training period that HIIT group train, the control mice remained in their cages.

### In vivo gene transfer in adult skeletal muscle

Tibialis anterior electroporation was performed as previously described [53]. Briefly, mice were anesthetized with 2-3% isoflurane. Hyaluronidase (H3884, Sigma) dissolved in sterile saline solution (0.36 mg/ml) was injected intramuscularly in tibialis anterior (TA) muscle, followed by 40 µg plasmid injection 1 hour later in re-anesthetized mice. Muscle electroporation was then performed by delivering 10 electrical pulses at an intensity of 100 V/cm, 10-ms pulse duration, 200-ms pulse interval using a caliper electrode (#45-0101 Caliper Electrode, BTX Caliper Electrodes, USA) connected to an ECM 830 BTX electroporator (BTX Harvard Apparatus). The p47-roGFP biosensor used to determine NOX2 activity was a kind gift from Dr. George G. Rodney [16].

### Redox Histology

Muscle freezing and sectioning was performed as previously reported [54]. In brief, TA muscles were dissected and embedded in optimum cutting temperature (OCT) medium from Tissue Tek, frozen in liquid nitrogen cooled isopentane and kept at −80 °C until processing. Redox histology was performed as previously described [55], Briefly, p47roGFP-transfected muscles were cut in 10 µm thickness followed by incubation in 50 μl of PBS containing 50 mM n-ethylmaleimide (NEM) for 10 min at 4°C. Sections were fixed in PBS-dissolved 4% paraformaldehyde (50 mM NEM) for 10 min, washed three times in PBS (5 min) and mounted in mounting medium (Vectashield, USA).

### Western Blotting

Western blotting was performed as previously described [12]. Briefly, ∼40 µg of quadriceps muscle and the whole soleus were lysed for 1 min at 30 Hz on a shaking bead-mill (TissueLyser II, Qiagen, Valencia, CA, USA) in ice-cold lysis buffer (0.05 mol/L Tris Base pH 7.4, 0.15 mol/L NaCl, 1 mmol/ L EDTA and EGTA, 0.05mol/L sodium flouride, 5 mmol/L sodium pyrophosphate, 2 mmol/L sodium orthovanadate, 1 mmol/L benzamidine, 0.5% protease inhibitor cocktail (P8340, Sigma Aldrich), and 1% NP-40). After rotating end-over-end for 30 min at 4°C, lysate supernatants were collected by centrifugation (18,327 g) for 20 min at 4°C. Lysate protein concentrations were determined using BSA standards (Pierce) and bicinchoninic acid assay reagents (Pierce). Total protein and phosphorylation levels of relevant proteins were determined by standard immunoblotting techniques, loading equal amounts of protein. The primary antibodies used; p-AMPK^Thr172^ (Cell Signaling Technology (CST)), #2535S), p-p38 MAPK^Thr180/Tyr182^ (CST, #9211), Hexokinase II (CST, #2867), GLUT4 p-ACC2 Ser^212^ (Millipore, 03-303), (ThermoFisher Scientific, PA-23052), Rac1 (BD Biosciences, #610650), NOX2 (Abcam, #Ab129068), Catalase (SCBT, sc-271803), SOD2 (Millipore, 06-984), TRX2 (SCBT, sc-50336), actin (CST, #4973) total p38 MAPK (CST, #9212), alpha2 AMPK (a gift from D. Grahame Hardie, University of Dundee), total ERK 1/2 (CST, #9102), TBC1D1^ser231^ (Millipore #07-2268), Hexokinase II (CST, #2867) MFN2 (#9482), and OPA1 (BD Biosciences #612606). A cocktail antibody (Abcam, #ab110413) was used as representative of the mitochondrial electron chain complexes (Oxphos). The optimal protein loading was pre-optimized to ensure measurements in the linear dynamic range for each antibody. Bands were visualized using a ChemiDoc imaging system (Bio-Rad, USA). Total protein staining (Coomassie) was used as a loading control rather than house-keeping proteins, as previously recommended [56]. Grayscale levels range for the blots shown was minimally and linearly adjusted across entire blots, as recommended by the American Society for Biochemistry and Molecular Biology.

### Single fiber immunostaining

Staining of single fibers was performed as previously described with slight modifications [30]. Briefly, ∼20 μg of quadricep muscles were fixed by immersion in ice-cooled 4% formaldehyde for 4h and long-term stored in 50% glycerol (diluted in PBS) at −20 °C. Muscles were teased into single fibers with fine forceps and transferred to immunobuffer (50 mM glycine, 0.25% bovine serum albumin (BSA), 0.03% saponin and 0.05% sodium azide in PBS) After isolation of a minimum of 30 muscle fibers were incubated overnight with an anti-COX4 antibody (#16056, Abcam, Cambridge, UK) in immunobuffer containing 0.5% saponin and, after 3 washes with immunobuffer, single muscle fibers were incubated for 2h with a secondary antibody conjugated with Alexa Fluor 488 (Invitrogen, UK). A negative control was performed with fibers not exposed to the primary antibody. The muscle fibers were mounted in Vectashield mounting medium.

### Imaging acquisition

All confocal images were collected using a 63x 1.4 NA oil immersion objective lens on an LSM 780 confocal microscope (Zeiss) driven by Zen 2011. Image acquisition was performed blinded. For the p47roGFP biosensor images, raw data of the 405- and 488-nm laser lines were exported to ImageJ as 16-bit TIFFs for further analysis. Data are represented as normalized fluorescence ratio (405/488 nm) and normalized to the WT untrained group.

### Statistical analyses

Results are shown as means ± S.E.M. Statistical testing was performed using *t*-tests, one-way or two-way (repeated measures when appropriate) ANOVA as applicable. Sidak’s post hoc test was performed when ANOVA revealed significant main effects. Statistical analyses were performed using GraphPad Prism 8.

## Author Contributions

CHO and TEJ designed research; CHO, LB, ZL, LAC, JRK, SHR, TEJ performed research; RH provided the *ncf1\** mice and intellectual input, CHO analyzed data; CHO and TEJ wrote the paper; all authors commented on the draft, TEJ Funding Acquisition.

## Acknowledgments

TEJ was supported by a Novo Nordisk Foundation Excellence project grant (#15182). CHO was supported by a CONICYT Ph.D. Scholarship. ZL was supported by a Chinese Scholarship Council Ph.D. stipend. JRK was supported by a Danish Diabetes Academy Ph.D. stipend. RH was supported by the Swedish Strategic Science Foundation (SSF). We thank Prof. Henriette Pilegaard for providing MFN2 and OPA1 antibodies. Imaging data were collected at the Center for Advanced Bioimaging and the Core Facility for Integrated Microscopy, University of Copenhagen, Denmark.

## Non-standard abbreviations

HIIE: high-intensity interval exercise
HIIT: high-intensity interval training
HK II: Hexokinase II
SOD2: superoxide dismutase 2
NOX2: NADPH oxidase 2
*ncf1\**: B10.Q. p47phox mutated PDH, pyruvate dehydrogenase
RER: respiratory exchange ratio
ROS: reactive oxygen species.

